# Polygenic routes lead to parallel altitudinal adaptation in *Heliosperma pusillum* (Caryophyllaceae)

**DOI:** 10.1101/2021.07.05.451094

**Authors:** Aglaia Szukala, Jessica Lovegrove-Walsh, Hirzi Luqman, Simone Fior, Thomas Wolfe, Božo Frajman, Peter Schönswetter, Ovidiu Paun

**Affiliations:** Department of Botany and Biodiversity Research, University of Vienna, Vienna, Austria; Vienna Graduate School of Population Genetics, Vienna, Austria; Department of Botany, University of Innsbruck, Innsbruck, Austria; Department of Environmental System Science, ETH Zürich, Zürich, Switzerland; Institute for Forest Entomology, Forest Pathology and Forest Protection, BOKU, Vienna, Austria

**Keywords:** Altitudinal adaptation, *Heliosperma pusillum*, Polygenic architecture, Parallel divergence, RNA-seq, Demography, Ecotypes

## Abstract

Understanding how organisms adapt to the environment is a major goal of modern biology. Parallel evolution - the independent evolution of similar phenotypes in different populations - provides a powerful framework to investigate the evolutionary potential of populations, the constraints of evolution, its repeatability and therefore its predictability. Here, we quantified the degree of gene expression and functional parallelism across replicated ecotype formation in *Heliosperma pusillum* (Caryophyllaceae), and gained insights into the architecture of adaptive traits. Population structure analyses and demographic modelling support a previously formulated hypothesis of parallel polytopic divergence of montane and alpine ecotypes. We detect a large proportion of differentially expressed genes (DEGs) underlying divergence within each replicate ecotype pair, with a strikingly low amount of shared DEGs across pairs. Functional enrichment of DEGs reveals that the traits affected by significant expression divergence are largely consistent across ecotype pairs, in strong contrast to the non-shared genetic basis. The remarkable redundancy of differential gene expression indicates a polygenic architecture for the diverged adaptive traits. We conclude that polygenic traits appear key to opening multiple routes for adaptation, widening the adaptive potential of organisms.

## Introduction

Independent instances of adaptation with similar phenotypic outcomes are powerful avenues for exploring the mechanisms and timescale of adaptation and divergence (Arendt and Reznick 2008; Turner et al. 2010; Agrawal 2017; Buckley et al. 2019; Knotek et al. 2020). A broad range of parallel to non-parallel genetic solutions can be causal to phenotypic similarity. Thus, evolutionary replicates converging to a similar phenotypic optimum offer insight into the constraints on evolution and help disentangle the nonrandom or more “predictable” actions of natural selection from confounding stochastic effects such as drift and demography (Lee and Coop 2019). In particular, repeated formation of conspecific ecotypes (Nosil et al. 2009; Nosil et al. 2017) are pivotal to enhancing our understanding of the processes leading to adaptation in response to a changing environment.

A number of studies have shown that parallelism at the genotype level can be driven by either standing genetic variation, possibly shared across lineages through pre- or post-divergence gene flow (Cooper et al. 2003; Colosimo et al. 2005; Jones et al. 2012; Soria-Carrasco et al. 2014; Van Belleghem et al. 2018; Alves et al. 2019; Thompson et al. 2019; Louis et al. 2020), or, more rarely, by recurrent *de novo* mutations with large phenotypic effects (Hoekstra et al. 2006; Chan et al. 2010; Zhen et al. 2012; Projecto-Garcia et al. 2013; Tan et al. 2020). These sources of adaptive variation produce phenotypic similarities via the same genetic locus, regardless of whether it was acquired independently or present in the ancestral gene pool (Stern 2013).

On the other hand, there is compelling evidence of phenotypic convergence resulting from non-parallel signatures of adaptation (Elmer et al. 2014; Yeaman et al. 2016; Rellstab et al. 2020), even among closely related populations (Wilkens and Strecker 2003; Steiner et al. 2009; Fischer et al. 2021) and replicated laboratory evolution (Barghi et al. 2019). A typical example is the convergent evolution of a lighter coat pigmentation in beach mouse populations of the Gulf of Mexico and the Atlantic Coasts driven by different mutations (Steiner et al. 2009).

Such cases suggest that evolutionary replicates can follow diverse non-parallel genetic routes and relatively few molecular constraints exist in the evolution of adaptive traits (Arendt and Reznick 2008; Losos 2011). The degree of parallelism during adaptation to similar selective pressures across taxa reveals that genomic signatures of adaptation are often redundant (Wilkens and Strecker 2003; Mandic et al. 2018; Fischer et al. 2021). The evolution of phenotypic similarity can involve highly heterogeneous routes depending on variation in gene flow, strength of selection, effective population size, demographic history, and extent of habitat differentiation, leading to different degrees of parallelism (MacPherson and Nuismer 2017; Yeaman et al. 2018). This complex range of processes including non-parallel to parallel trajectories have also been described using the more comprehensive term *continuum of (non)parallel evolution* (Stuart et al. 2017; Bolnick et al. 2018).

Recently, a quantitative genetics view of the process of adaptation has gained attention among evolutionary biologists (Barghi et al. 2020), complementing existing models on adaptation via selective sweeps. Accordingly, selection can act on different combinations of loci, each of small effect, leading to shifts in the trait mean through changes in multiple loci within the same molecular pathway (Hermisson and Pennings 2017; Höllinger et al. 2019). Thus, key features of polygenic adaptation are that different combinations of adaptive alleles can contribute to the selected phenotype (Barghi et al. 2020) and that the genetic basis of adaptive traits is fluid, due to the limited and potentially short-lived contribution of individual genetic loci to the phenotype (Yeaman 2015). This genetic redundancy (Goldstein and Holsinger 1992; Nowak et al. 1997; Láruson et al. 2020) can lead to non-parallel genomic changes in populations evolving under the same selective pressure. Footprints of selection acting on polygenic traits have been detected in a wide range of study systems, such as in fish (Therkildsen et al. 2019) and in cacao plants (Hämälä et al. 2020), potentially fostering convergent adaptive responses and phenotypes during independent divergence events (Lim et al. 2019; Rougeux et al. 2019; Hämälä et al. 2020).

A current major challenge is predicting adaptive responses of populations and species to environmental change. Despite several advances, it is still unclear which adaptive signatures are expected to be consistent across evolutionary replicates, especially when selection acts on complex traits. Important aspects to investigate are the architecture of adaptive traits (simple/monogenic, oligogenic or polygenic) and the repeatability of genetic responses in independent instances of adaptation (Yeaman et al. 2018). A polygenic architecture may facilitate alternative pathways leading to the same phenotypic innovation, diminishing the probability of parallel evolution at the genotype level, but likely enhancing the adaptive potential of populations at the phenotypic level (Boyle et al. 2017). To date, we observe a steady increase of plant studies addressing (non-)parallel evolution both at the genotype and phenotypic level (e.g. Roda et al. 2013; Yeaman et al. 2016; Trucchi et al. 2017; Cai et al. 2019; Konečná et al. 2019; Rellstab et al. 2020; Tan et al. 2020; Bohutínská et al. 2021; James, Arenas-Castro et al. 2021; James, Wilkinson et al. 2021). Still more attention should be given to specifically assessing parallelism in light of the idea of genetic redundancy that have been emphasized over the past few years.

Altitudinal ecotypes of *Heliosperma pusillum* s.l. (Waldst. & Kit.) Rchb. (Caryophyllaceae) offer a system to study this process. In the Alps, this species includes an alpine ecotype (1,400–2,300 m above sea level) widely distributed across the mountain ranges of southern and central Europe, and a montane ecotype (500–1,300 m) endemic to the south-eastern Alps (Fig. 1a). The latter was previously described from scattered localities as *H. veselskyi* Janka, but the two ecotypes are highly interfertile (Bertel, Hülber, et al. 2016) and isolation-by-distance analyses confirmed their conspecificity (Trucchi et al. 2017). While the alpine ecotype has a relatively continuous distribution in moist screes above the timberline, the montane ecotype forms small populations (typically < 100 individuals) below overhanging rocks.

**Figure 1.**
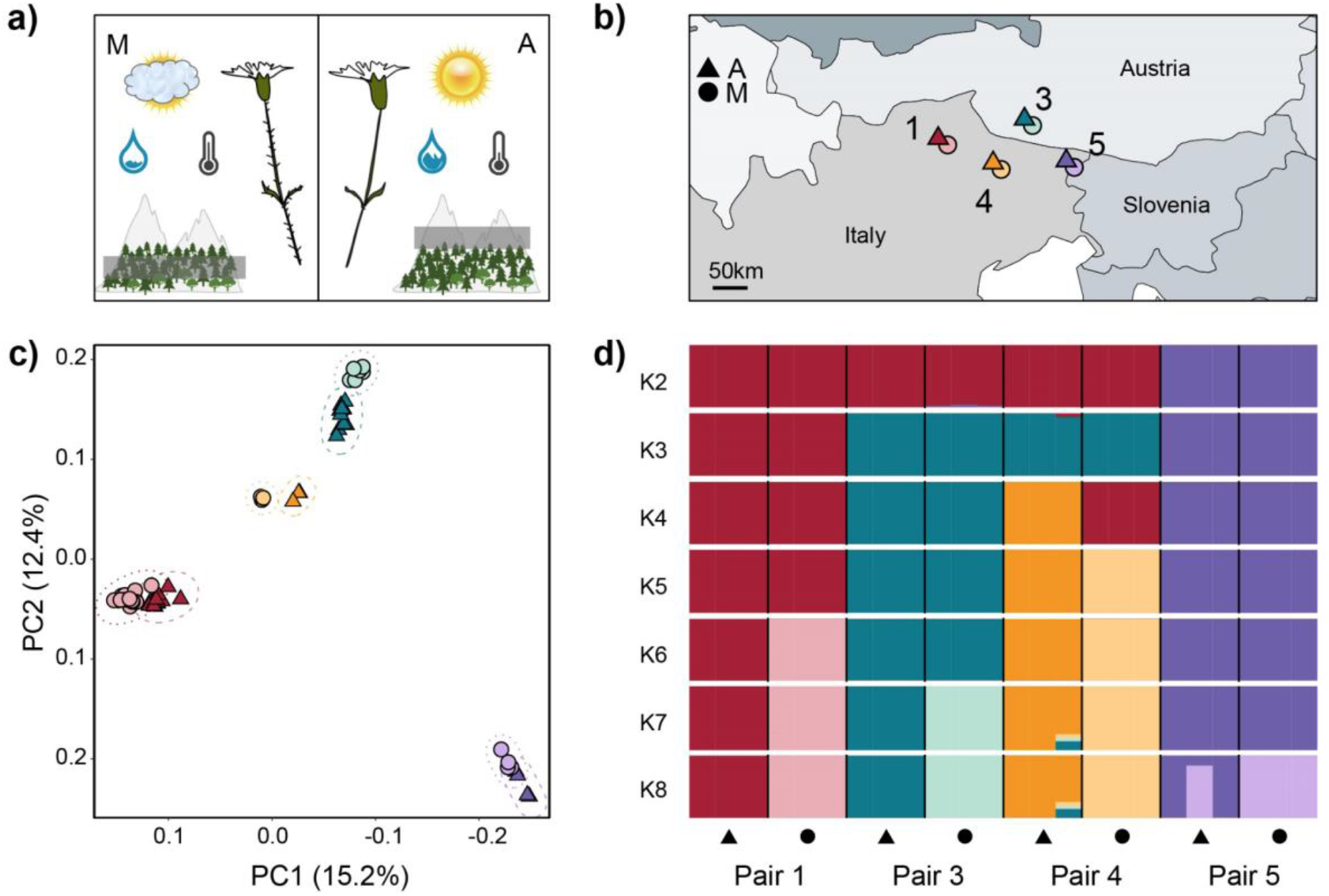
Study system, sampling setup, and genetic variation among four montane (M, circles) - alpine (A, triangles) ecotype pairs of *Heliosperma pusillum*. Color coding of populations is consistent across panels. The numbering of the ecotype pairs is consistent with previous work (Bertel et al. 2018). **(a)** Graphic description of the main ecological and morphological differences between the ecotypes. **(b)** Geographic map showing the location of the analyzed populations in the southeastern Alps. **(c)** Clustering of individuals along the first two vectors of a principal component analysis. **(d)** Bar plot showing the assignment of individuals to the clusters identified by NgsAdmix for K = 2 through 8.

Previous work (Bertel, Buchner, et al. 2016; Bertel et al. 2018) reported substantial abiotic differences between the habitats preferred by the two ecotypes. For example, differences in average temperature (montane: warm vs. alpine: cold), temperature amplitude, the degree of humidity (montane: dry vs. alpine: humid) and light availability (montane: shade vs. alpine: full sunlight) were found between the two altitudinal sites. Moreover, metagenomics (Trucchi et al. 2017) showed evidence of distinct microbial communities in the respective phyllospheres. The two ecotypes also differ significantly in their physiological response to light and humidity conditions in a common garden (Bertel, Buchner, et al. 2016). Finally, the montane ecotype is covered by a dense glandular indumentum, which is absent in the alpine populations (Frajman and Oxelman 2007; Bertel et al. 2017).

Both ecotypes show higher fitness at their native sites in reciprocal transplantation experiments (Bertel et al. 2018), confirming an adaptive component to their divergence. Common garden experiments across multiple generations further rejected the hypothesis of a solely plastic response shaping the phenotypic divergence observed (Bertel et al. 2017). Most importantly, population structure analyses based on genome-wide SNPs derived from restriction site-associated DNA (RAD-seq) markers (Trucchi et al. 2017) supported a scenario of five parallel divergence events across the six investigated ecotype pairs. Hereafter, we use the term “ecotype pairs” to indicate single instances of divergence between alpine and montane ecotypes across their range of co-occurrence.

The combination of ecological, morphological, and demographic features outlined above renders *H. pusillum* a well-suited system to investigate the mechanisms driving local recurrent altitudinal adaptation in the Alps. Here, we quantify the magnitude of gene expression and functional parallelism across ecotype pairs, by means of RNA-seq analyses of plants grown in a common garden. We also investigate the independent evolution of ecotype pairs more in depth than previously. More specifically, this study asks (1) how shared are gene expression differences between ecotypes among evolutionary replicates or, in other words, is the adaptation to elevation driven by expression changes in specific genes or in different genes affecting similar traits; (2) how shared is the functional divergence encoded by differentially expressed genes (DEGs) among evolutionary replicates; and (3) do we find consistent signatures of selection on coding sequence variation across evolutionary replicates?

## Materials and methods

### Reference genome assembly and annotation

We assembled *de novo* a draft genome using short and long read technologies for an alpine individual of *H. pusillum* that descended from population 1, from a selfed line over three generations. DNA for long reads was extracted from etiolated tissue after keeping the plant for one week under no light conditions. DNA was extracted from leaves using a CTAB protocol adapted from Cota-Sánchez et al. (2006). Illumina libraries were prepared with IlluminaTruSeq DNA PCR-free kits (Illumina) and sequenced as 150 bp paired-end reads on Illumina HiSeq X Ten by Macrogen Inc. (Korea). PacBio library preparation and sequencing of four SMRT cells on a Sequel I instrument was done at the sequencing facility of the Vienna BioCenter Core Facilities.

MaSuRCA v.3.2.5 (Zimin et al. 2013) was used to perform a hybrid assembly using 192.3 Gb (ca 148×) Illumina paired-end reads and 14.9 Gb (ca 11.5×) PacBio single-molecule long reads. The assembled genome was structurally annotated *ab initio* using Augustus (Stanke et al. 2006) and GeneMark-ET (Lomsadze et al. 2014), as implemented in BRAKER1 v.2.1.0 (Hoff et al. 2016) with the options *--softmasking=1 --filterOutShort*. Mapped RNA-seq data from three different samples was used to improve *de novo* gene finding.

A transcriptome was assembled using Trinity v.2.4.0 (Haas et al. 2013) to be used in MAKER-P v.2.31.10 (Campbell et al. 2014) annotation as expressed sequence tag (EST). We used as additional evidence the transcriptome of the closely related *Silene vulgaris* (Sloan et al. 2011). The annotation was further improved during the MAKER-P analyses by supplying gene models identified using BRAKER1, and by masking a custom repeat library generated using RepeatModeler v.1.0.11 (http://www.repeatmasker.org/RepeatModeler/). Gene models identified by both BRAKER1 and MAKER-P were functionally annotated using Blast2GO (Götz et al. 2008; Haas et al. 2013). BUSCO v.2.0 (Simão et al. 2015) was used for quality assessment of the assembled genome and annotated gene models using as reference the embryophyta_odb10 dataset.

### Sampling, RNA library preparation and sequencing

Our main aim was to test the repeatability of the molecular patterns and functions that distinguish the alpine from the montane ecotype in different ecotype pairs. To achieve this goal, we performed DE analyses on 24 plants grown in common garden settings at the Botanical Garden of the University of Innsbruck, Austria. Wild seeds were collected from four alpine/montane ecotype pairs in the south-eastern Alps (Fig. 1b, Table S1). The numbering of localities is consistent with that used in Bertel et al. (2018) and the acronyms corresponding to Trucchi et al. (2017) are added in Table S1. All seeds were set to germination on the same day and the seedlings were grown in uniform conditions. One week before RNA fixation, the plants were brought to a climate chamber (Percival PGC6L set to 16 h 25 °C three lamps/8 h 15 °C no lamps). Then, fresh stalk-leaf material, sampled at a similar developmental stage for all individuals, was fixed in RNAlater in the same morning and kept at −80 °C until extraction. Total RNA was extracted from ca 90 mg leaves using the mirVana miRNA Isolation Kit (Ambion) following the manufacturer’s instructions. Residual DNA has been digested with the RNase-Free DNase Set (Qiagen); the abundant ribosomal RNA was depleted by using the Ribo-Zero rRNA Removal Kit (Illumina). RNA was then quantified with a NanoDrop2000 spectrophotometer (Thermo Scientific), and quality assessed using a 2100 Bioanalyzer (Agilent). Strand-specific libraries were prepared with the NEBNext Ultra Directional RNA Library Prep Kit for Illumina (New England Biolabs). Indexed, individual RNA-seq libraries were sequenced with single-end reads (100 bp) on 11 lanes of Illumina HiSeq 2500 at the NGS Facility at the Vienna BioCenter Core Facilities (VBCF; https://www.viennabiocenter.org/). Two samples (A1a and A4b) were sequenced with paired-end reads (150 bp) with the initial aim of assembling reference transcriptomes.

To identify genetic variants under selection we extended the sampling by including 41 additional transcriptomes of individuals from ecotype pairs 1 and 3 (Fig. 1b) grown in a transplantation experiment (Szukala A et al. unpublished data; Table S1). The procedure used to prepare the RNA-seq libraries was the same as described above, except that the indexed, individual libraries have been sequenced with single-end reads (100 bp) on Illumina NovaSeq S1 on 2 lanes at the Vienna BioCenter Core Facilities.

### Genetic diversity and structure

RNA-seq data was demultiplexed using BamIndexDecoder v.1.03 (http://wtsi-npg.github.io/illumina2bam/#BamIndexDecoder) and raw sequencing reads were cleaned to remove adaptors and quality filtered using trimmomatic v.0.36 (Bolger et al. 2014). Individual reads were aligned to the reference genome using STAR v.2.6.0c (Dobin et al. 2013). Mapped files were sorted according to the mapping position and duplicates were marked and removed using Picard v.2.9.2 (https://broadinstitute.github.io/picard/). The individual bam files were further processed using GATK v.3.7.0 function IndelRealigner to locally improve read alignments around indels. Subsequently, we used a pipeline implemented in ANGSD v.0.931 (Korneliussen et al. 2014) to estimate genotype likelihoods. The latter might be more reliable than genotype calling for low coverage segments, in particular when handling data with strongly varying sequencing depth among regions and individuals such as RNA-seq. Briefly, ANGSD was run to compute posterior probabilities for the three possible genotypes at each variant locus (considering only bi-allelic SNPs), taking into account the observed allelic state in each read, the sequencing depth and the Phred-scaled quality scores. ANGSD was run with the options *-GL 2 -doMajorMinor 1 -doMaf 1 -SNP_pval 2e-6 -minMapQ 20 -minQ 20 -minInd 12 -minMaf 0.045 -doGlf 2*. A significant portion of RNA-seq data includes protein coding regions expected to be under selection. To investigate genetic structure and demography the dataset was further filtered to keep genetic variants at four-fold degenerate (FFD) sites using the Bioconductor package VariantAnnotation in R (Obenchain et al. 2014).

A covariance matrix computed from the genotype likelihoods of FFD variants at unlinked positions (i.e., one per 10 Kb windows) was used for principal component analysis using PCAngsd v.0.99 (Meisner and Albrechtsen 2018). To test for admixture, we run NgsAdmix (Skotte et al. 2013) on genotype likelihoods at FFD unlinked sites. The number of clusters tested for the admixture analysis ranged from K = 1 to K = 9. The seed for initializing the EM algorithm was set to values ranging from 10 to 50 to test for convergence. Finally, the K best explaining the variance observed in the data was evaluated using the Evanno method (Evanno et al. 2005) in CLUMPAK (http://clumpak.tau.ac.il/bestK.html). Result plotting was performed using R v.3.5.2.

For each population we estimated the average global Watterson’s theta (θw) and average pairwise nucleotide diversity (π), whereas to test for departures from mutation/drift equilibrium we computed Tajima’s D (Tajima 1989). Global average estimates of these statistics were computed using ANGSD and custom bash scripts, implementing a sliding window approach with windows of 50 Kb and a step of size 10 Kb.

We estimated between-population differentiation as *F*_ST_ for all pairs of populations at high and low elevation respectively, as well as for pairs of ecotypes across localities. *F*_ST_ statistics were carried out in ANGSD using the folded joint site frequency spectra (jSFS) for all population pairs as summary statistics. Given that no suitable outgroup sequence was available, the ancestral state was unknown. As a consequence, we observed a deviation from the expected SFS for some populations (i.e. a high frequency of sites with fixed alternate alleles) when polarizing toward the major allele throughout the alpine populations. Therefore, we produced site allele frequency likelihoods using ANGSD settings *-dosaf 1 -GL 2 -minQ 20 -P 8 -skipTriallelic 1 -doMajorMinor 1 -anc reference.genome.fasta*, limiting the analysis to the set of FFD sites using the *-sites* option. Finally, we used the *-fold* option to fold the spectra when using realSFS (for further analyses in ANGSD), and using a custom R script to fold the spectra into fastsimcoal2 format (for coalescent simulations in fastsimcoal2).

### Testing alternative demographic scenarios

We performed coalescent simulations to differentiate between two different possible explanations behind the patterns of genetic structure observed. One possible scenario implies multiple, polytopic divergence events between the ecotypes, whether or not gene flow was involved. Another possibility is that the two ecotypes diverged only once, whereas subsequent gene flow between ecotypes in each pair could have homogenized their genetic background. Therefore, we tested two contrasting topologies for each combination of two ecotype pairs (Fig. 2): one model assuming a single origin (1-origin) of each ecotype, and one assuming independent between-ecotype divergence across geographic localities (2-origins). Additionally, for each topology two scenarios were evaluated: one in absence of migration between populations (strict isolation, SI) and one with continuous migration between demes (isolation with migration, IM). In line with the results from the population structure analyses our expectation was to find higher migration rates between ecotypes within each ecotype pair (solid lines in Fig. 2).

**Figure 2.**
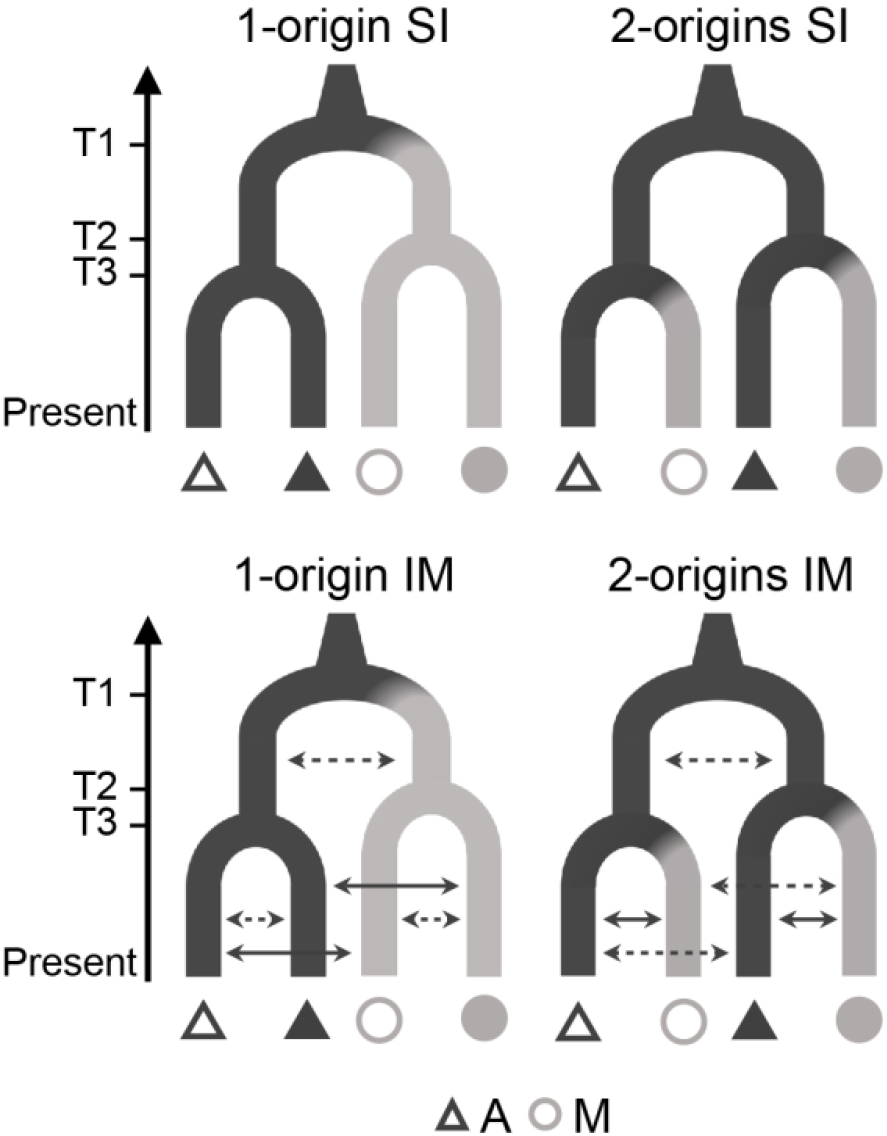
Alternative topologies tested using Fastsimcoal2 for all combinations of two ecotype pairs. Strict isolation (SI, upper panels) and isolation with migration (IM, lower panels) were modeled. Solid arrows in the IM models indicate higher migration rates expected between ecotypes at each locality according to population structure results. Divergence times T2 and T3 were allowed to vary (i.e., T2 > T3 but also T3 > T2 were modeled), whereas T1 was always the oldest event. Triangles and circles represent populations of the alpine (A) and the montane (M) ecotype, respectively. Filled and empty symbols represent different ecotype pairs.

We evaluated which demographic scenario (1-origin vs 2-origins) explains our data using fastSimcoal2 v.2.6.0.3 (Excoffier et al. 2013). We tested four populations at a time, i.e. with two ecotype pairs in each simulation, using for each analysis the jSFS for all six combinations of populations as summary statistics. For all models we let the algorithm estimate the effective population size (N), the mutation rate (μ) and the time of each split (T1, T2 and T3, Fig. 2). Although N, μ and the time of split between ecotypes in each pair have been previously estimated by Trucchi et al (2017), we started with broad search ranges for the parameters to not constrain a priori the model. The final priors of the simulations were set for a mutation rate between 1e-8 and 1e-10, the effective population size between 50 and 50,000 (alpine populations) and 50 and 5,000 (montane populations), and for the time of each split between 1,000 and 100,000 generations ago. We forced T1 to predate T2 and T3, and performed separate simulations setting T2 > T3 and T3 > T2, respectively. For the models including gene flow, migration rate (m) between any pair of demes was initially set to a range between 10e-10 and two.

The generation time in *H. pusillum* was reported to be 1 year (Flatscher et al. 2012; Trucchi et al. 2017). While most populations in the montane zone flower during the first year after germination, this is not the case in the alpine environment, where plants usually start to flower in the second year after germination. Therefore, 1 year is most likely an underestimation of the intergeneration interval, which is more realistically around 3 years. While this parameter does not affect the overall results in terms of topology, it should be considered carefully in terms of divergence times between ecotypes that were previously hypothesized to be post-glacial (Flatscher et al. 2012; Trucchi et al. 2017).

FastSimcoal2 was run excluding monomorphic sites (*-0* option). We performed 200,000 simulations and ran up to 50 optimizations (ECM) cycles to estimate the parameters. To find the global optimum of the best combination of parameter estimates, we performed 60 replicates of each simulation run. The MaxEstLhood is the maximum estimated likelihood across all replicate runs, while the MaxObsLhood is the maximum possible value for the likelihood if there was a perfect fit of the expected to the observed site frequency spectrum. We report the difference between these two estimates (ΔL) for each model and ΔAIC scores (i.e., the difference between the AIC for the best possible model and the tested model) to compare models with different numbers of parameters. Finally, the parameter estimations of the best run were used to simulate the expected jSFS and test the goodness of fit of the topology plus parameter estimates to the observed data.

### Differential gene expression analysis

Only unique read alignments were considered to produce a table of counts using FeatureCounts v.1.6.3 (Liao et al. 2014) with the option *-t gene* to count reads mapping to gene features. DE analyses were performed using the Bioconductor package EdgeR v.3.24.3 (Robinson et al. 2010). The count matrix was filtered, keeping only genes with mean counts per million (cpm) higher than 1. Data normalization to account for library depth and RNA composition was performed using the weighted trimmed mean of M-values (TMM) method. The estimateDisp() function of edgeR was used to estimate the trended dispersion coefficients across all expressed tags by supplying a design matrix with ecotype pair and ecotype information for each sample. We implemented a generalized linear model (glm) to find gene expression differences between low and high elevation ecotypes by taking into account the effects of the covariates ecotype and ecotype pair on gene expression. A likelihood ratio test (lrt) was used to test for DE genes between ecotypes in each pair. The level of significance was adjusted using Benjamini-Hochberg correction of p-values to account for multiple testing (threshold of FDR < 0.05). The statistical significance of the overlaps between lists of DEGs was tested using a hypergeometric test implemented in the Bioconductor package SuperExactTest (Wang et al. 2015) and the number of genes retained after trimming low counts as background. Finally, to compare the repeatability of gene usage in DEGs to the neutral expectation and to the repeatability of selection outliers detected (see below), we computed the Jaccard index for any two ecotype pairs and the C-hypergeometric score metric that was specifically developed with the scope of comparing repeatability of the evolutionary process across multiple lineages (Yeaman et al. 2018).

### Functional interpretation of DEG

We performed separate gene ontology (GO) enrichment analyses for the lists of DEGs of each ecotype pair and gave special attention to functions that were shared among lists of DEG. We also performed similar GO terms enrichments after excluding any DEGs shared between at least two ecotype pairs. This additional analysis was performed to clarify if sets of fully non-shared DEGs would result in similar enriched functions. Fisher test statistics implemented in the Bioconductor package topGO v.2.34.0 (https://bioconductor.org/packages/release/bioc/html/topGO.html) were run with the algorithm “weight01” to test for over-representation of specific functions conditioned on neighbouring terms. Multiple testing correction of p-values (FDR correction) was applied and significance was assessed below a threshold of 0.05. DEGs were also explicitly searched for protein coding genes and transcription factors underlying the formation of trichomes and visually checked using R.

### Detection of multilocus gene expression variation

To detect gene expression changes underlying adaptive traits with a strongly polygenic basis we performed a conditioned (partial) redundancy analysis (cRDA) of the gene expression data using the R package vegan v.2.5-6 (Oksanen et al. 2019). The cRDA approach is well suited to identify groups of genes showing expression changes that covary with the “ecotype” variable while controlling for population structure (Bourret et al. 2014; Forester et al. 2018). As a table of response variables in the cRDA we used the cpm matrix after filtering using a mean cpm higher than 1 as in the DE analysis. First, the cRDA includes a multiple regression step of gene expression on the explanatory variable “ecotype”. In our case, the RDA was conditioned to remove the effects of the geographic ecotype pair using the formula “~ ecotype + Condition(pair)”. In the second step, a principal component analysis (PCA) of the fitted values from the multiple regression is performed to produce canonical axes, based on which an ordination in the space of the explanatory variable is performed. The first axis of the cRDA therefore shows the variance explained by the constrained variable “ecotype”, while the second axis is the first component of the PCA nested into the RDA, representing the main axis of unconstrained variance. The significance of the cRDA was tested with ANOVA and 1,000 permutations. Each gene was assigned a cRDA score that is a measure of the degree of association between the expression level of a gene and the variable “ecotype”. Outliers were defined as genes with scores above the significance thresholds of ± 2 and, respectively, ± 2.6 standard deviations from the mean score of the constrained axis, corresponding to p-value thresholds of 0.05 and 0.01, respectively.

### SNPs calling and detection of selection outliers

To detect outlier genetic variants potentially under divergent selection during ecotype adaptation to different elevations, we computed per locus *F*_ST_ based on the sfs of the genotype likelihoods computed in ANGSD. Selection outliers analyses were carried out on ecotype pairs 1 and 3, for which we had a minimum of 10 individuals in each population analyzed. To account for low coverage values in DEGs, a site would be retained if a minimum low coverage of four would be found in at least seven individuals. Consequently, ANGSD was run with the options *-dosaf 1 -GL 2 -minQ 20 -MinMapQ30 -skipTriallelic 1 -doMajorMinor 1 -doCounts 1 -setMinDepthInd 4 -minInd 7 -setMaxDepthInd 150*. We then computed the sfs using the *-fold 1* option and ran the ANGSD script *realSFS* with the option *-whichFst 1* to compute the Bhatia et al. (2013) *F*_ST_ estimator by gene following the procedure described at https://github.com/ANGSD/angsd/issues/239. We then defined as *F*_ST_ outliers those loci falling in the top 5% of the *F*_ST_ distribution. To understand if DEGs carry stronger signatures of selection than other genes, we compared the *F*_ST_ distribution of 1,000 randomly selected genes to the *F*_ST_ distribution of DEGs and tested the difference in mean using a permutation test. Finally, we computed the Jaccard index and C-hypergeometric score (Yeaman et al. 2018) to compare repeatability in selection outliers to the repeatability in usage of DEGs.

## Results

### Reference genome assembly and annotation

Our hybrid *de novo* genome assembly recovered a total length of 1.21 Gb of scaffolds corresponding to 93% of the estimated genome size (1C = 1.3 pg; Temsch et al. 2010). The draft *H. pusillum* genome v.1.0 is split into 75,439 scaffolds with an N50 size of 41,616 bp. RepeatModeler identified 1,021 repeat families making up roughly 71% of the recovered genome. This high proportion of repetitive elements aligns well with observations in other plant genomes.

Structural annotations identified 25,661 protein-coding genes with an average length of 4,570 bp (Fig. S2a and b). All protein-coding genes were found on 8,632 scaffolds that belong to the longest tail of the contig length distribution (Fig. S2c). Nevertheless, we also observe in our assembly comparatively long contigs that do not contain any gene models (Fig. S2c). Of the total set of genes, 17,009 could be functionally annotated (Götz et al. 2008; Haas et al. 2013). We evaluated the completeness of the genome assembly by searching our gene models against the BUSCO v.3 embryophyta dataset (Simão et al. 2015). When running BUSCO on the annotated mRNA, a total of 82.4% of the set of single-copy conserved BUSCO genes were found. A BUSCO search on the part of the genome remaining after hard masking genes, could still identify 9.6%conserved BUSCO orthologs within ‘non-genic’ regions. This Whole Genome Shotgun project has been deposited at DDBJ/ENA/GenBank under the accession JAIUZE000000000.

### Genetic diversity and structure

Two alpine individuals of pair 3 (A3b and A3c, Table S1) were found to be highly introgressed with genes from the alpine population of pair 4 (Fig. S1a), and have been discarded from subsequent genetic analyses, retaining a total of 63 individuals for further analyses based on SNPs. This dataset was also used to test the hypothesis of parallel ecotype divergence in *H. pusillum* suggested by Trucchi et al (2017).

To explore population diversity and structure we filtered a dataset of 7,107 putatively neutral variants at unlinked FFD sites from 63 individuals representing the four ecotype pairs (Fig. 1b, Table S1). Within-population allelic diversity (average pairwise nucleotide diversity, π, and Watterson’s theta, θ_w_), Tajima’s D, as well as *F*_ST_, are reported in Table S2. Average π showed similar values across alpine and montane populations, ranging from π_A5_ = 0.140 ± 0.12 to π_A1_ = 0.173 ± 0.12 in the alpine ecotype, and from π_M3_ = 0.143 ± 0.12 to π_M5_ = 0.171 ± 0.13 in the montane. Watterson’s theta ranged from θ_w-A5_ = 0.130 ± 0.11 to θ_w-A4_ = 0.136 ± 0.12 and from θ_w-M3_ = 0.116 ± 0.08 to θ_w-M5_ = 0.157 ± 0.11 in the alpine and montane ecotype, respectively. We did not observe a clear alpine versus montane distinction of within-population allelic diversity. Global Tajima’s D estimates were always positive, but close to 0 (Table S2, Fig. S3), suggesting that these populations are within neutral-equilibrium expectations, and that both alpine and montane populations were not affected by major changes in population size in the recent past.

Averaged pairwise *F*_ST_ tended to be slightly higher between montane than between alpine populations (weighted *F*_ST_ = 0.28-0.56 for alpine, and weighted *F*_ST_ = 0.39-0.52 for montane; Table S2). Between-ecotype *F*_ST_ was lower than *F*_ST_ between pairs, except in the case of pair 4 (weighted *F*_ST_ = 0.48), consistent with overall high expression differentiation between ecotypes in this pair as described below.

We further investigated the population structure with principal component analyses (PCA) and an admixture plot, both based on genotype likelihoods computed in ANGSD v.0.931 (Korneliussen et al. 2014). In the PCA (Fig. 1c) the analyzed populations cluster by geography, in line with previous results (Trucchi et al. 2017). The first component (15.2% of explained variance, Fig. 1c) shows a clear east-west separation of the ecotype pairs. The second component (12.4% of explained variance, Fig. 1c) places ecotype pair 5 closer to pair 1 and most distant from pair 3 showing a north-south separation.

We performed two rounds of population structure inference using NgsAdmix v.32 (Skotte et al. 2013), to test the effects of uneven sample size on the inferred clusters. We compared the results inferred using the set of 63 accessions to those inferred when randomly subsampling all populations to three individuals (i.e., the minimum number of individuals per population in our dataset). With uneven sampling, we observed that the individuals from populations with reduced sampling size (i.e., ecotype pair 4) tended to be assigned to populations of higher sampling density (Fig. S1b), an otherwise known problem affecting population structure analyses (Puechmaille 2016; Meirmans 2019). Consistent with the clustering observed in the PCA, pair 5 was first separated from the other pairs (K = 2, Fig. 1d). The best three Ks, as evaluated using the Evanno method (Evanno et al. 2005), were 2, 3 and 7, in this order, confirming an enhanced separation of pair 5 from the rest, while the two ecotypes in this pair are the least diverged (K = 7, Fig. 1d), consistent with a lower degree of expression differentiation in this pair (Fig. 3).

**Figure 3.**
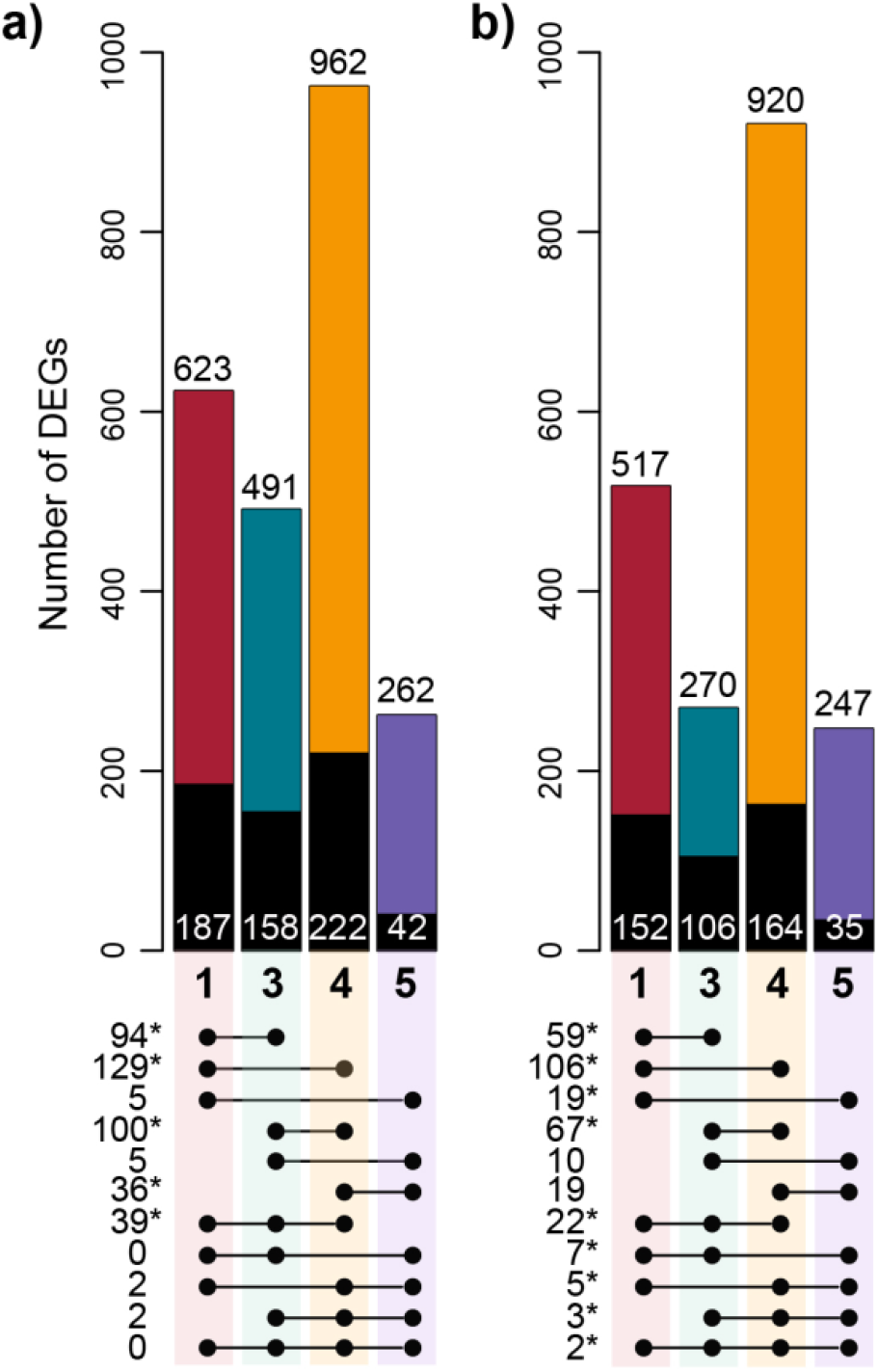
Differentially expressed genes (DEGs) at each ecotype pair show low overlap across different pairs. Colors represent the ecotype pair as in Fig. 1. Histograms show the number of DEGs (FDR < 0.05) underexpressed **(a)** and overexpressed **(b)** in the montane ecotype compared to the alpine in each pair. Colored and black areas of the bars show the amount of DEGs unique to each ecotype pair and, respectively, shared with at least one other pair. Numbers reported on top of the bars show the total amount of DEGs between ecotypes per pair and category. Numbers on the black areas show the amount of DEGs shared with at least one other pair. Linked dots below bars show the amount of shared DEGs between two, three or four pairs. Stars indicate that the overlap is significantly higher than chance expectations (hypergeometric test, p < 0.01).

### Demographic model selection, parallelism and gene flow

Delta Akaike information criteria (ΔAIC) for each demographic model tested in fastSimcoal2 are summarized in Table S3a and c. In the absence of gene flow (SI models), our simulations consistently showed that the 2-origins topologies are preferred over the 1-origin hypotheses. However, IM models (i.e., allowing gene flow) always achieved a higher likelihood than SI models (Table S3a). The 2-origins IM scenario again achieved a better likelihood in five out of six ecotype pairs comparisons. The 1-origin IM model was preferred for pairs 3-4. For each parameter we took as a final estimate the 95% confidence intervals CI of the ten best model estimates. The CI of the times of divergence and effective population size (Ne) from the best model estimates are reported in Table S3b. We computed migration rate estimates for each model including both directions of migration for all combinations of ecotype populations from two pairs (Table S3d). We found migration rates to be very low across all comparisons and scenarios tested (upper limit of the CI always below 0.015); generally they were estimated to be lower between different ecotype pairs than between ecotypes in each pair (Table S3d).

### Patterns of differential gene expression between ecotypes

We analyzed gene expression in a common garden to identify genes with divergent expression between ecotypes, as these are hypothesized to underlie phenotypic differentiation and adaptation to different altitudinal niches. After trimming genes with low expression across samples we retained a dataset of 16,389 genes on which we performed DE analyses.

A major proportion of DEGs were found to be unique to each pair (colored area of the bars in Fig. 3a and b). This pattern was particularly enhanced in pair 5, in which ca 85% of DEGs were not shared with other pairs, while *ca* 70%, 65% and 80% of DEGs were unique to pairs 1, 3 and 4, respectively (Fig. 3, Fig. S4). Although the overlap of DEGs was significantly higher than chance expectations (p < 0.01) for several comparisons, our analyses recovered an overall low number of shared DEGs. In contrast to expectations, we found across all ecotype pairs that only two and zero genes were consistently over- and under-expressed in the montane compared to the alpine ecotype, respectively. Consistently, Jaccard similarity indexes computed for any two ecotype pairs were very low, lying between 0.005 and 0.09 (Table S4). Given the null expectation that any gene in our trimmed dataset could contribute to ecotype divergence (i.e., background set including 16,389 genes), C-hypergeometric scores across all pairs were equal to 8.46 and 9.16 for genes under- and overexpressed in the montane ecotype compared to the alpine.

The number of DEGs varied relatively widely across ecotype pairs. DEGs were almost four times higher in pair 4 (highest degree of expression differentiation) compared to pair 5 (lowest degree of expression differentiation), while the difference in DEGs was less pronounced between pairs 1 and 3. This result is consistent with the PCA of normalized read counts (Fig. S5a) and the multidimensional scaling plot of gene expression (Fig. S5b). The relative degree of expression differentiation between ecotypes at different geographic localities is also consistent with their degree of genetic differentiation (*F*_ST_, Table S2). The second component of the PCA of gene expression (13.8% of variance explained, Fig. S5a), as well as the second dimension of logFC of the multidimensional scaling analysis (Fig. S5b), tend to separate the two ecotypes. Interestingly, gene expression appears more uniform across the montane accessions compared to the alpine ones, even if the overall expression divergence between different populations was not significantly different between ecotypes (Wilcoxon signed rank test p = 0.56; Fig. S6 and Table S5).

### Parallel multilocus gene expression variation

We performed a cRDA of gene expression to elucidate if a different analytical framework would provide more power to detect common genes with opposite expression patterns between ecotypes across all evolutionary replicates. Redundancy analysis is thought to be a good approach to detect changes between conditions (in our case, ecotypes), even when such differences are subtle and possibly masked by other factors (Forester et al. 2018).

We found that 1.8% of total expression variation was explained by divergence between montane and alpine ecotypes across all ecotype pairs (Fig. 4), consistent with the low overlap of DEGs across evolutionary replicates. Also consistent with the low number of shared DEGs, the ANOVA test of the full model was not significant (F = 1.39, p = 0.18), confirming that most expression differences between ecotypes in our dataset do not follow consistent routes across ecotype pairs. We further searched for cRDA outliers to identify genes with consistent, albeit subtle, changes in expression across ecotypes. The transcript score was transformed into a z-score following Wolfe et al. (2021) with a distribution ranging from −3.55 to 3.43 (Fig. S7). We identified 115 genes at a significance level p < 0.01 (2.6 SD), and 739 at a significance p < 0.05 (2 SD) with an outlier expression between the two ecotypes that was consistent across all pairs. Overlaps with DEGs identified in edgeR are reported in Fig. S8.

**Figure 4.**
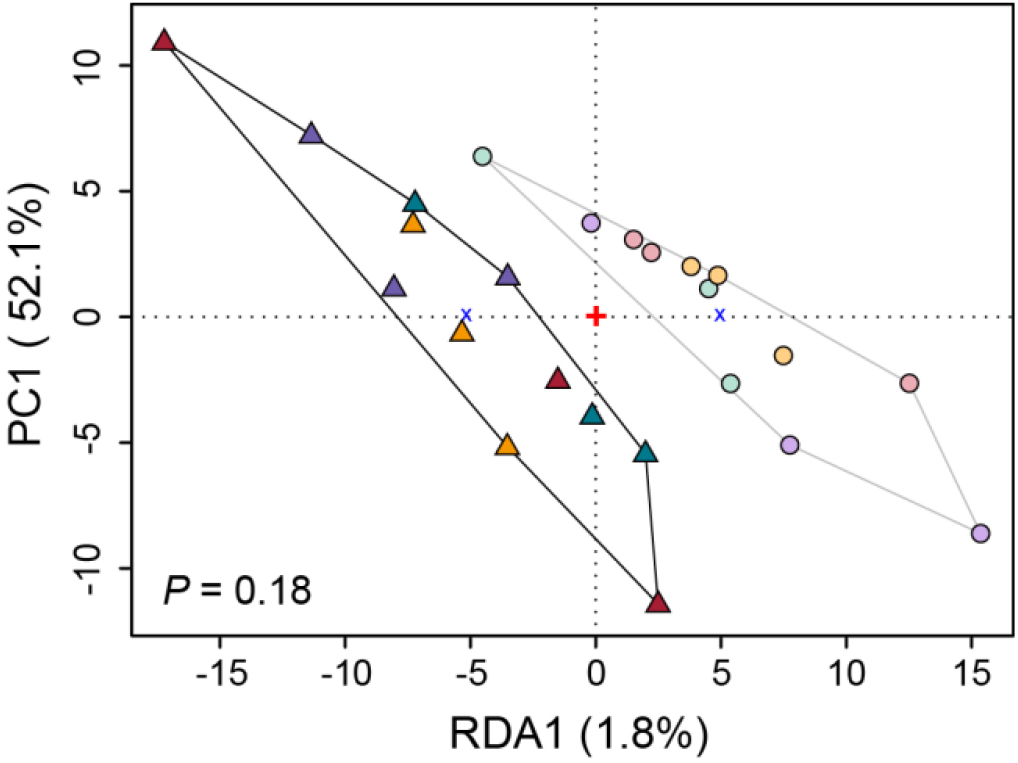
Expression divergence between accessions of the alpine and the montane ecotypes captured with conditioned redundancy analysis (cRDA). Colors represent the populations as in Fig. 1. Triangles and circles represent alpine (A) and montane (M) individuals, respectively, whereas the black and grey lines delimited clusters correspond to the alpine and the montane ecotypes, respectively. The ANOVA test of the full model was not significant (p = 0.18), confirming that most expression differences between ecotypes in our dataset do not follow consistent routes across ecotype pairs.

### Ecological and biological significance of DEGs

In stark contrast to the low overlap at the level of individual genes affected by DE, we observed evidence of convergence in the enriched biological functions across DEG lists of each ecotype pair. To allow easier interpretation, we exemplify in Fig. 5 a subset of the significantly enriched GO terms that can be easily related to the ecological and morphological ecotype divergence. Enrichments among all DEGs (Fig. 5a, Table S6a), but also after excluding shared DEGs (Fig. 5b, Table S6b) are reported. We observed that GO terms enriched (adjusted p < 0.05) in genes that were differentially expressed without exclusion of shared DEGs included trichome development, light and cold response, drought response including regulation of stomatal activity,, responses to biotic stress and plant growth (Fig. 5a, Table S6a). These enrichments appeared to be largely consistent among the different ecotype pairs, even after excluding the shared DEGs (Fig. 5b, Table S6b). The z-score indicated that the GO terms related to trichome development were represented by genes that tended to be overexpressed in the montane ecotype (Fig. 5a and b), while the overall degree of over- and underexpression of genes underlying other convergent GO terms across pairs varied depending on the specific function of the genes affecting the respective molecular pathway. We also analyzed enriched biological processes in cRDA gene outliers (Table S7), since these genes possibly underlie biologically and ecologically relevant adaptive traits. Consistent with the DE results, cRDA outlier genes were significantly enriched for defense responses, including jasmonic and salicylic acid related pathways, as well as response to light, cold, ozone and water deprivation.

**Figure 5.**
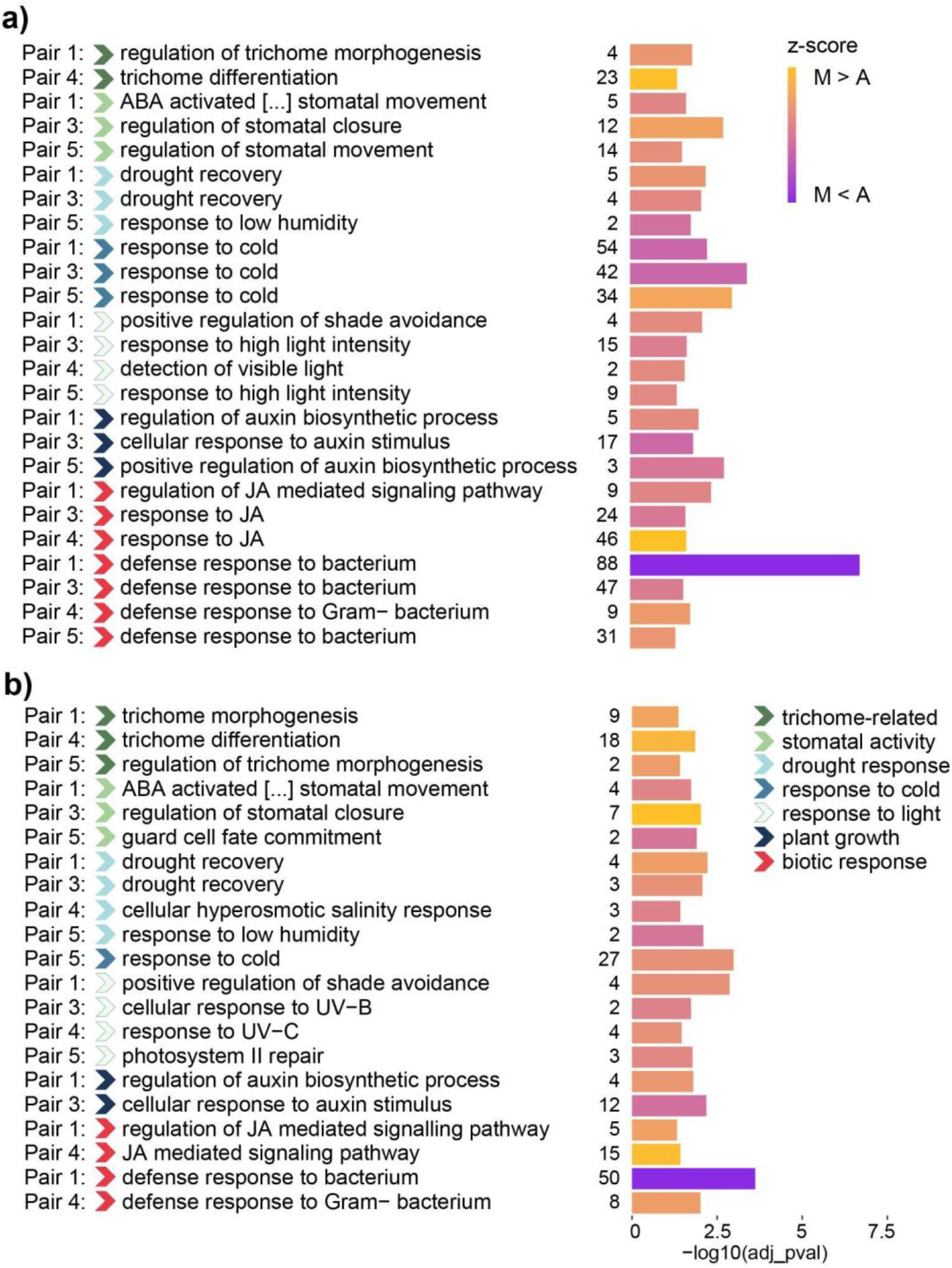
Functional enrichment of differentially expressed genes (DEGs), showing that across ecotype pairs similar biological processes appear linked to adaptation to the different elevations. GO terms enrichment including all DEGs **(a)** and excluding shared DEGs **(b)**. The ecotype pair in which a certain term is found to be enriched is specified on the left side of the plots. The broad category to which the GO terms pertain is indicated with colored arrows, according to the legend. The size of the bars shows the adjusted significance of the enriched GO terms (Fisher’s test). Numbers left of the bars show the number of DEGs underlying the corresponding GO term. The z-score (color scale of the bars) was computed based on the log fold-change of gene expression, whereas positive and negative values show over- and underexpression in the montane ecotype respectively. ABA, abscisic acid; JA, jasmonic acid; UV, ultraviolet radiation.

In the GO enrichment analysis of the cRDA outliers we did not find significantly enriched GO terms related to trichome development. Consistently, the genes underlying this trait identified in DE analyses were largely not shared by different ecotype pairs. We observed that some genes known to be involved in trichome formation in *A.thaliana* and found to be expressed in our transcriptomes were significantly differentially expressed in some of the ecotype pairs but not in others, or showed consistent changes in expression between ecotypes even if not significant after FDR correction (examples shown in Fig. 6). For instance, the gene IBR3, a Indole-3-butyric acid response gene, known to promote hair elongation (Strader et al. 2010; Velasquez et al. 2016) was always overexpressed in the montane ecotype as compared to the alpine (Fig. 6). This same gene was also significantly differentially expressed in three out of four ecotype pairs in previous DEG analyses before correction of p-values for multiple testing (Fig. 6).

**Figure 6.**
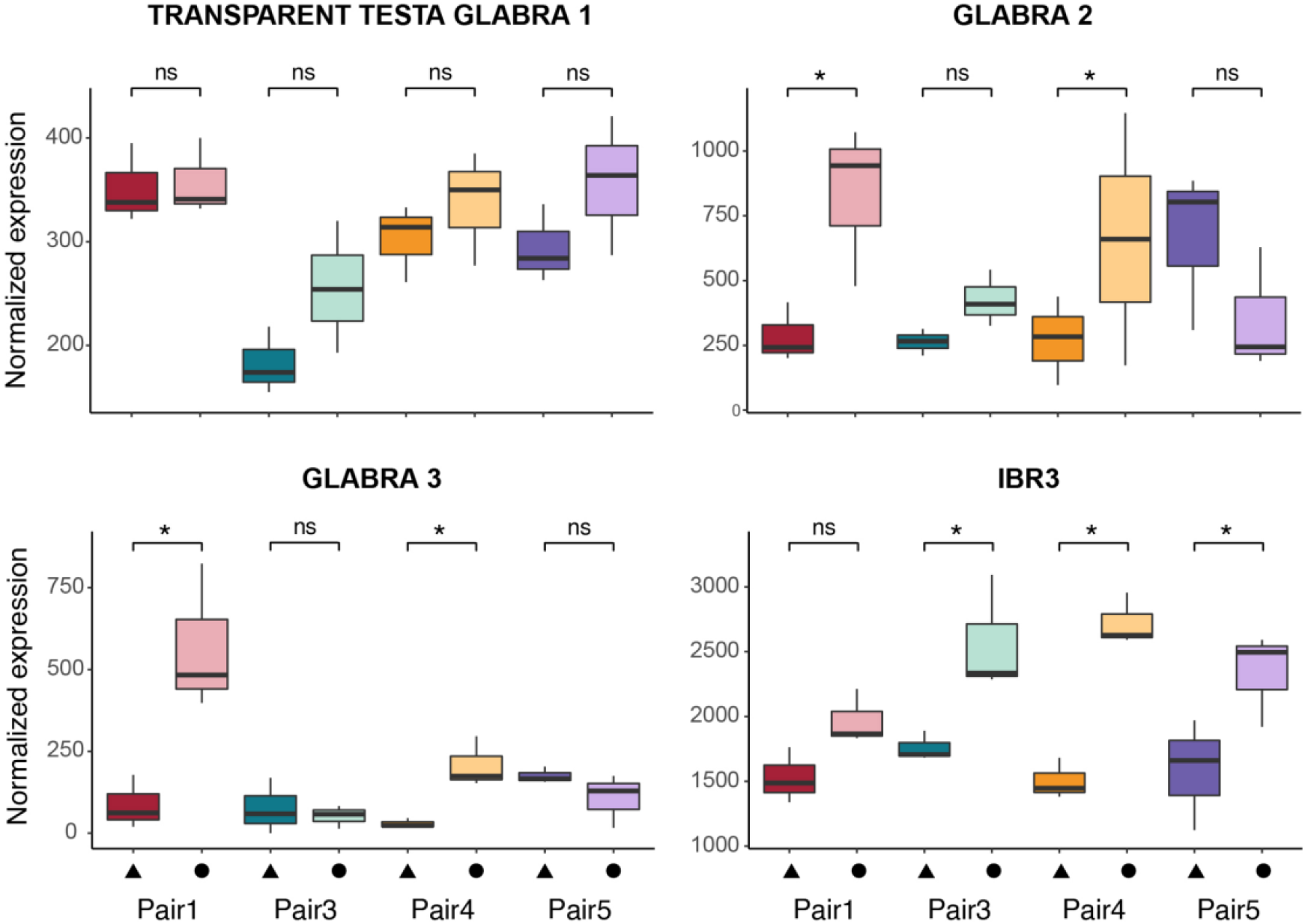
Examples of expression of genes known to be related to trichome formation and elongation in plants. Colors represent the populations as in Fig. 1. Triangles and circles represent populations of the alpine (A) and the montane (M) ecotype, respectively. Stars indicate significant differential expression (p < 0.05) before false discovery rate (FDR) correction. Non-significant differences are marked with ns.

### (Non-)Shared adaptive outlier loci

To identify possible candidate genes under divergent selection in independent divergence events, we searched for coding genomic regions with pronounced allelic divergence between ecotypes in pairs 1 and 3. We excluded ecotype pairs 4 and 5 from this analysis because of the low number of individuals available from these populations.

Two sets of 3,300 and 2,811 genes were retained in pair 1 and 3, respectively, for *F*_ST_ analyses with 2,766 genes shared by both pairs. We found that the *F*_ST_ distribution of DEGs in each pair did not significantly differ from the *F*_ST_ distribution of 1,000 randomly selected genes (Fig. S9, permutation test p = 0.4 in both pair 1 and 3), suggesting that the identified DEGs were not positioned in regions under stronger selection than other protein coding regions. We detected 165 and 141 *F*_ST_ outlier genes in pair 1 and 3, respectively. Eighteen genes containing outlier SNPs were shared by both pairs, a number significantly higher than expected by chance (p = 0.001). The lower Jaccard index recovered in selection outliers (Jaccard index = 0.0003) compared to DEGs of these ecotype pairs (Jaccard index equal to 0.092 and 0.081 for genes under- or overexpressed in the montane compared to the alpine of the pairs 1 and 3, Table S4) indicates that the similarity of selection outliers is even less pronounced than the similarity of DEGs. We recovered a C-hypergeometric score equal to 4.2, which confirms that the shared *F*_ST_ outlier genes are less distant from the null expectation than the overlap of DEGs. Functional annotations of the 18 shared genes containing outlier SNPs are reported in Table S8. Among those candidate genes, we found genes involved in defense response (At1g53570, At3g18100), ion channel and transport activity (At5g57940, At3g25520, At1g34220), and regulation of transcription and translation (At3g18100, At3g25520, At1g18540). Ten and seven *F*_ST_ outlier genes were also differentially expressed in pair 1 and 3, respectively, but not shared by both pairs..

## Discussion

Parallel evolution has long been recognised as a powerful process to study adaptation, overcoming intrinsic limitations of studies on natural populations that often miss replication (Elmer and Meyer 2011). In this work, we aimed to investigate the genetic basis of adaptation to different elevations in the plant *Heliosperma pusillum*. We asked in particular to what extent different ecotype pairs show signatures of parallel evolution in this system.

Our genetic structure analyses and coalescence-based demographic modelling were in line with a scenario of parallel, polytopic ecotype divergence, as suggested previously by a marked dissimilarity of the genomic landscape of differentiation between ecotype pairs revealed by RAD-seq data (Trucchi et al. 2017). In our demographic investigations, parallel divergence always obtained greater support under a strict isolation model. Still, models including low amounts of gene flow were shown to be more likely. Additionally, in one comparison (i.e., including ecotype pairs 3 and 4) the single origin IM scenario aligned more closely with the data than the two origins IM. This result is consistent with greater co-ancestry observed for these two pairs with respect to other comparisons (Fig. 1c and d). Nevertheless, the estimates of migration rates between different ecotype pairs were overall extremely low (i.e., always lower than 1.2e-03), indicating that each ecotype pair diverged in isolation from other pairs, even when it is not straightforward to distinguish between the different models (i.e. 1-origin vs 2-origins) in the case of pairs 3 and 4.

Our results from selection scans showed that only few diverged genes, likely under selection during adaptation to different elevations, were shared between the two ecotype pairs analyzed (i.e., pair 1 and 3), while over 87% of putatively adaptive loci were unique to each pair. This high degree of unique outliers, consistent with RADseq results from a previous investigation (Trucchi et al. 2017), supports a scenario of mainly independent evolutionary history of different ecotype pairs. However, we cannot exclude the possibility that a few shared loci, likely from standing genetic variation, might have played a role in shaping the ecotype divergence of different evolutionary replicates in our system.

Global Tajima’s D estimates were close to 0, suggesting that the recent past of all these populations was not affected by major bottlenecks or population expansions. Consistently, within-population diversity was similar across montane and alpine ecotypes, likely reflecting ancestral variation before altitudinal divergence. Due to the low number of individuals available for ecotype pairs 4 (three individuals per ecotype) and 5 (four individuals per ecotype), these estimates should be considered with caution. However, previous work using an RNA-seq-derived dataset of synonymous variants similar to ours (Fraïsse et al. 2018) showed that model selection based on the joint site frequency spectrum is robust to the numbers of individuals and loci. Nevertheless, future analyses should aim for enlarged sampling sizes.

We further asked how consistent across divergence events are the molecular processes underlying ecotype formation. We screened the expression profiles of four ecotype pairs grown in a common garden to shed light on the genetic architecture of the adaptive traits involved in parallel adaptation to divergent elevations, as well as to warmer/dry vs. colder/humid conditions. Our analyses showed that gene expression changes between ecotypes is largely genetically determined, and not a plastic response due to environmental differences. However, we found strikingly few DEGs shared across all four ecotype pairs, with most DEGs unique to one ecotype pair, suggesting that convergent phenotypes do not consistently rely on changes in expression of specific genes. Interestingly, montane populations were shown to be morphologically more diverged among each other than alpine populations, despite the similarity of ecological conditions across localities in both the montane and alpine niche (Bertel et al. 2018). Therefore, both morphological disparity, as well as different DEGs implicated in differentiation across lineages might reflect differing functional strategies to adapt to the montane/alpine environment.

The low sharedness of DEGs was most strongly driven by ecotype pair 5, which we also showed to bear a lower degree of shared ancestry with the other pairs in the genetic structure analyses (Fig. 1c-d). Given that ecotype pair 5 is the most eastern in terms of geographic distribution, it can be hypothesized that this pair represents a more distinct lineage, as break zones in the distribution of genetic diversity and distribution of biota have been identified to the West of this area of the Alps (Thiel-Egenter et al. 2011). This pair was shown also to be the earliest diverging among the four lineages included here (Trucchi et al. 2017), and this locality lies closest to the margin of the last glacial maximum (LGM) ice sheet. Following the retreat of the ice sheet, it is likely that this area could have been colonized first, whereas the ancestors of other ecotype pairs likely needed more time to migrate northwards prior to the onset of divergence. An alternative explanation might involve two different LGM refugia for pair 5 and the other three pairs. Our sampling was not appropriate to further test hypotheses of biogeographic nature. Even so, our results suggest that parallel evolution is analyzed at different levels of coancestry in our dataset. This implies that parallel signatures of ecotype evolution can decrease significantly, even within a relatively small geographic range. This view is in line with previous findings of unexpectedly heterogeneous differentiation between freshwater and marine sticklebacks across the globe, including more distant lineages (Fang et al. 2020).

Despite the low parallelism in gene activity, we identified across the ecotype pairs a high reproducibility of the biological processes related to ecological (i.e., different water and light availability, temperature and biotic stress) and morphological (i.e. glandular trichomes absence/presence) divergence at the two elevations. Functional enrichment of responses to biotic stress are consistent with the biotic divergence between the two habitat types, featuring distinct microbiomes (Trucchi et al. 2017) and accompanying vegetation (Bertel et al. 2018). The dichotomy of convergence in enriched GO terms, but a low amount of shared DEGs, indicates that different redundant genes likely concur to shape similar phenotypic differentiation, as expected under polygenic adaptation (Barghi et al. 2020). Shared genes containing selection outliers were involved in partly similar biological processes as those affected by DEGs, albeit noting that they may not be directly the targets of selection. Nevertheless, we found that shared selection outliers include regulatory elements of transcription, such as the MYB4R1 (gene At3g18100) transcription factor, and it is therefore possible that such *trans* regulatory elements under divergent selection cause at least part of the expression divergence observed. A largely *trans* control of expression divergence is consistent with our results that show that DEGs (together with their *cis* regulatory regions) do not generally reside within regions of high differentiation (i.e., *F*_ST_) between the ecotypes.

The presence (montane ecotype) or absence (alpine ecotype) of multicellular glandular hairs on the plants represents a striking morphological difference in our system. Trichome formation has been studied extensively in Brassicaceae, especially in *Arabidopsis*, where this trait is controlled by a relatively simple regulatory pathway shared across the family (Hülskamp et al. 1994; Hülskamp 2004; Hilscher et al. 2009; Pesch and Hülskamp 2009; Tominaga-Wada et al. 2011; Chopra et al. 2019). Still, a certain degree of genetic redundancy has been shown to underlie trichome formation in *Arabidopsis* (Khosla et al. 2014). Studies on other plant lineages, such as cotton (Machado et al. 2009), snapdragons (Tan et al. 2020), *Artemisia* (Shi et al. 2018) and tomato (Chang et al. 2018), highlighted that the genetic basis of multicellular glandular trichomes formation does not always involve the same loci as in *Arabidopsis*. Trichome formation outside of the Brassicaceae family likely involves convergent changes in different genetic components (Serna and Martin 2006; Tan et al. 2020) and has been reported to be initiated even as an epigenetic response to herbivory in *Mimulus guttatus* (Scoville et al. 2011).

We expected to find evidence of specific genes controlling trichome development in our transcriptome dataset. Indeed, we did observe a change in regulation of particular genes underlying trichome formation and elongation pathways across ecotype pairs. Interestingly, these genes were not-shared by different ecotype pairs, which was unexpected given the relatively simple genetic architecture of this trait in *A. thaliana*. Also, key genes known to underlie hair initiation in *A. thaliana*, or elongation and malformation in other plant species, were differentially expressed in some ecotype pairs, but not in all of them.

Analyses of replicated evolution in laboratory experiments on bacteria (Cooper et al. 2003; Fong et al. 2005), yeast (Nguyen Ba et al. 2019) and *Drosophila* (Barghi et al. 2019) have provided insights about adaptation, showing that redundant trajectories can lead to the same phenotypic optimum, when selection acts on polygenic traits. In line with other studies on diverse organisms including whitefish (Rougeux et al. 2019), hummingbirds (Lim et al. 2019), snails (Ravinet et al. 2016) and frogs (Sun et al. 2018), our results suggest that convergent phenotypes can be achieved via changes in different genes affecting the same molecular pathway and, ultimately, adaptive traits, and that this polygenic basis might facilitate repeated adaptation to different elevations via alternative routes.

In conclusion, this study adds evidence to recent findings showing that polygenic traits and genetic redundancy open multiple threads for adaptation, providing the substrate for reproducible outcomes in convergent divergence events. Future studies using transcriptomics as well as genomic approaches should focus on genotype-by-environment interactions, e.g., in reciprocal transplantation experiments, to further deepen our understanding of the process of adaptation in *H. pusillum*.

## Acknowledgements

This work was financially supported by the Austrian Science Fund (FWF) through the doctoral programme (DK) grant W1225-B20 to a faculty team, including O.P., and through grant Y661-B16 to O.P. We thank Nicholas Barton, Andrew Clark, Virginie Courtier-Orgogozo, Joachim Hermisson, Thibault Leroy, Magnus Nordborg, John Parsch, Christian Schlötterer, Emiliano Trucchi, Daniele Filiault and Alex Widmer for insightful comments and feedback. We thank Juliane Baar, Marie Huber, Carles Ferré Ortega and Daniela Paun for their support during laboratory work and data acquisition, Marianne Magauer and Daniela Pirkebner for producing the selfed line, as well as Martina Imhiavan and Daniel Schlorhaufer from the Botanical Garden of the University of Innsbruck for the cultivation of the plants. A permit to conduct the presented research activities was granted by the Parco Naturale Dolomiti Friulane (no. 1943); for the Austrian federal state Tirol no such permit was necessary.

## Data accessibility

All raw read data was uploaded to NCBI and can be found under Bioproject ID PRJNA760819. The new *Heliosperma pusillum* genome assembly v1.0 is available from GenBank (BioProject ID PRJNA739571, accession number JAIUZE000000000). Scripts used to perform the analyses can be found under this gitHub repository https://github.com/aglaszuk/Polygenic_Adaptation_Heliosperma/. More specifically, the raw table of read counts used for differential expression analysis can be found at https://github.com/aglaszuk/Polygenic_Adaptation_Heliosperma/tree/main/02_DifferentialExpression/data, while the lists of differentially expressed genes at https://github.com/aglaszuk/Polygenic_Adaptation_Heliosperma/tree/main/02_DifferentialExpression/results

## Author contributions

Study conceived and designed by OP, BF and PS. Sampling was performed by BF and PS. Laboratory work conducted by AS, JLW and OP. Bioinformatics and statistical analyses were conducted by AS and OP, with feedback from HL, SF and TW. Interpretation of the results was undertaken by all authors. The manuscript was first drafted by AS, and was revised and approved by all authors.

