## Supplementary figures for "Polygenic routes lead to parallel altitudinal adaptation in *Heliosperma pusillum* (Caryophyllaceae)": SI_12.21.pdf

**Figure S1. (a)** NgsAdmix results (K = 3 and 4) for ecotype populations in locality 3 and 4 showing that individuals A3b and A3c (marked on the figure) are highly introgressed from population A4. **(b)** NgsAdmix results (K = 2 through 8) for the ecotype pairs using the dataset including 63 accessions, i.e. without random downsampling of each population to the same number of individuals (compare to Fig. 1d in the main text). The colors represent different gene pools.

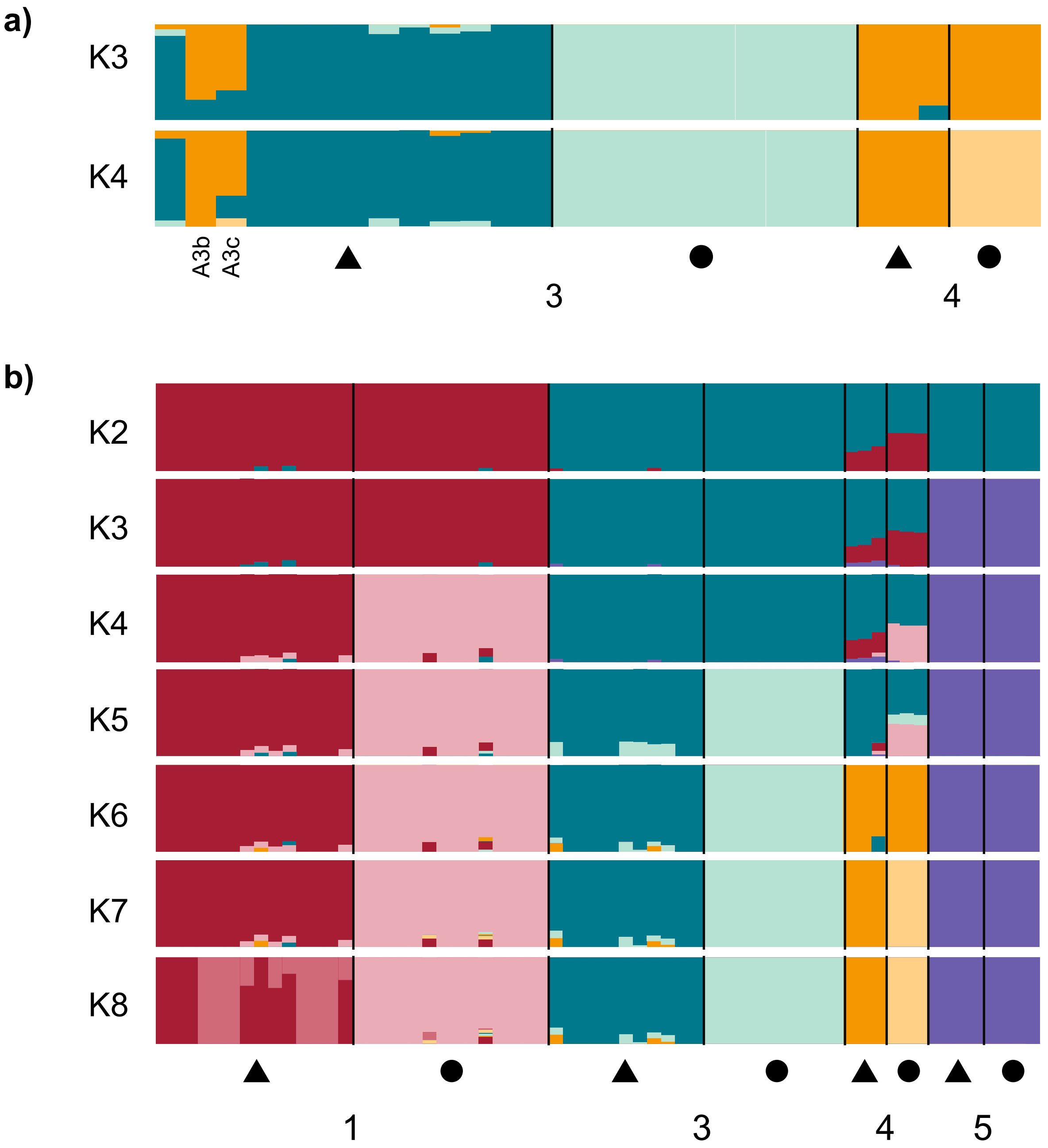

**Figure S2. (a)** Gene length distribution of the *Heliosperma* genome assembly. **(b)** Distribution of the amount of exons per gene. **(c)** Contig length distribution. Highlighted in darker grey color is the proportion of contigs containing genes.

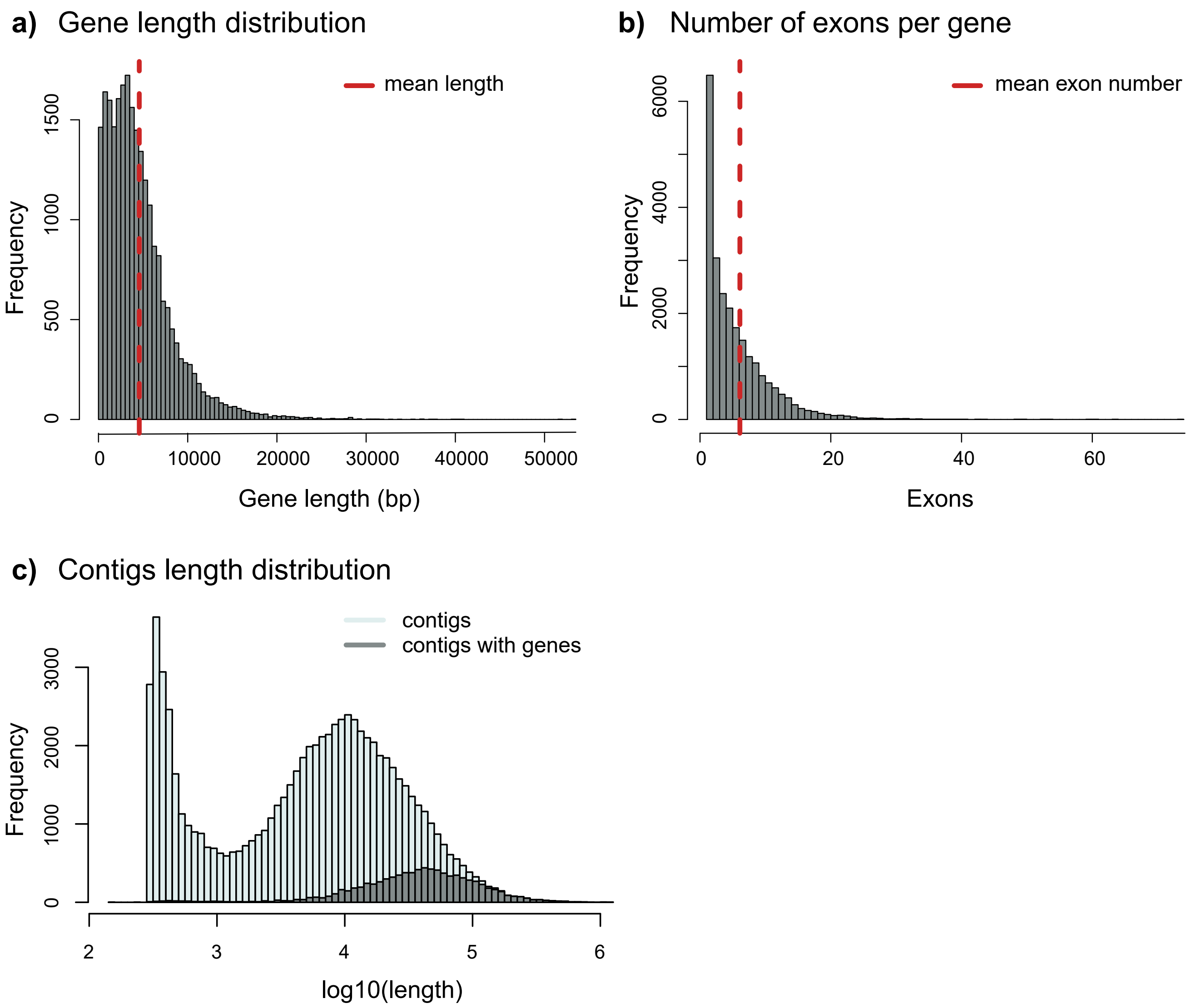

**Figure S3.** Population-wise distribution of Tajima's D estimates computed per window of size 50Kb with a step of 10Kb. Colors represent the populations as in Fig. 1 in the main text. A - alpine populations; M - montane populations.

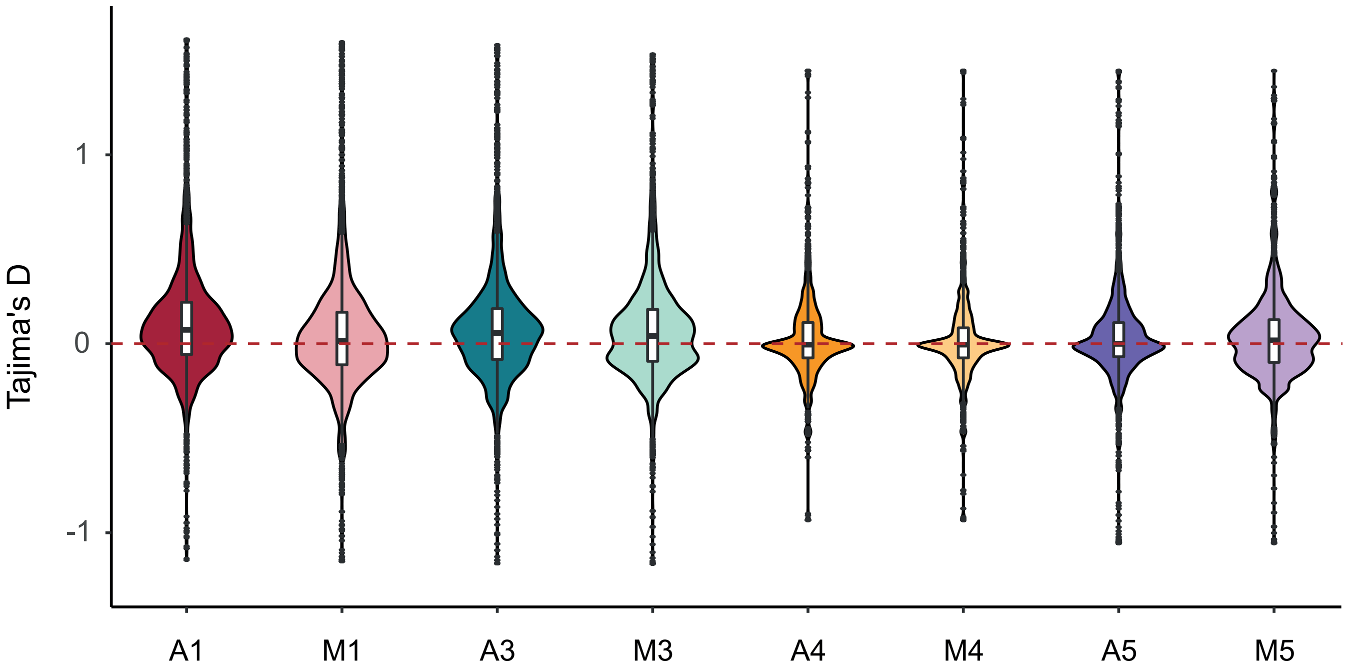

**Figure S4.** Scatterplots of differentially expressed genes between ecotypes, showing ecotype pairs comparisons and the amount of shared DEG (red dots).

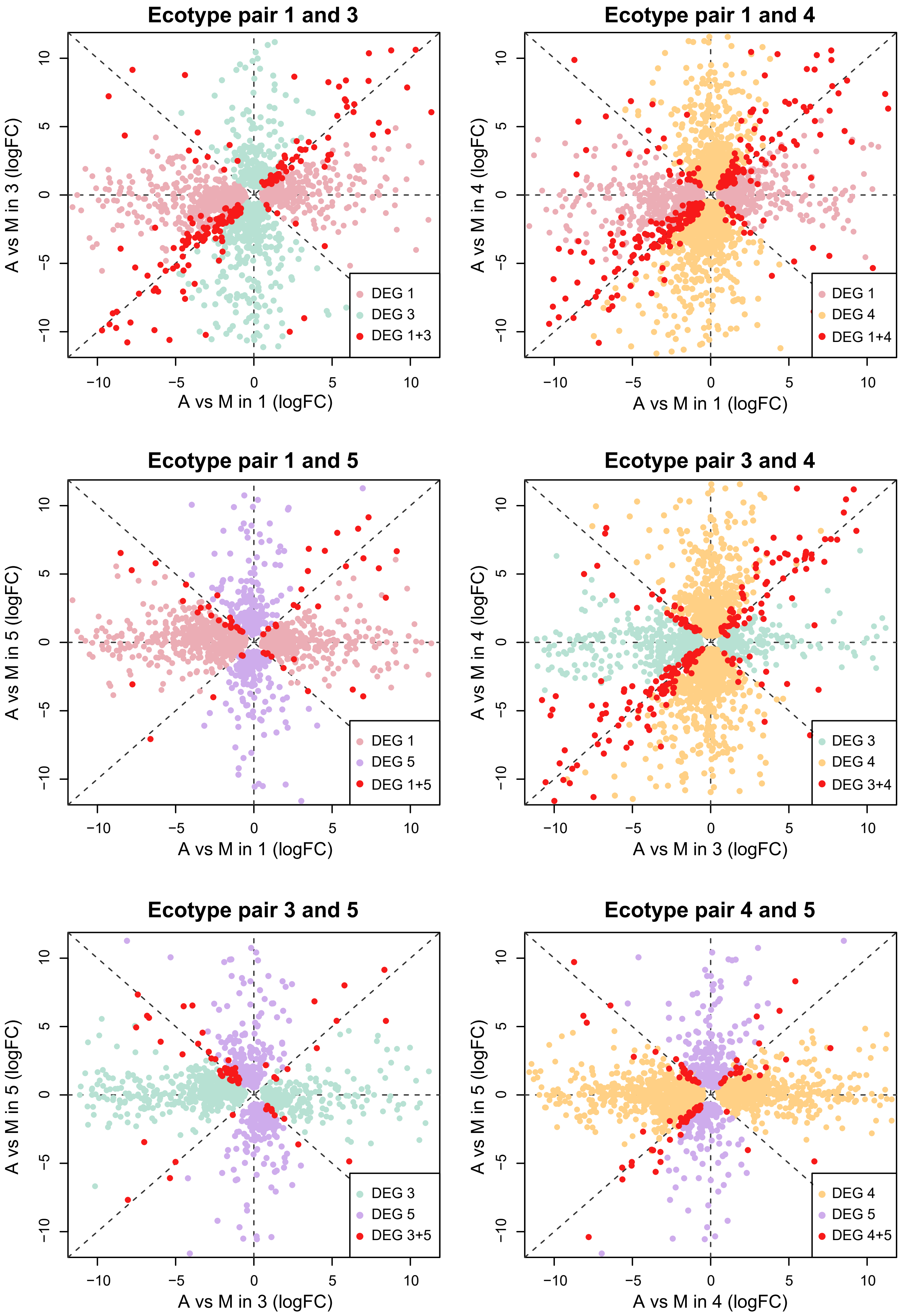

**Figure S5. (a)** PCA (dimensions 1 to 3) based on normalized gene expression counts. **(b)** Multidimensional scaling plots (dimensions 1 to 3) of logFC of gene expression counts after normalization. Colors represent populations as in Fig.1b. Circles represent montane individuals; triangles show alpine accessions.

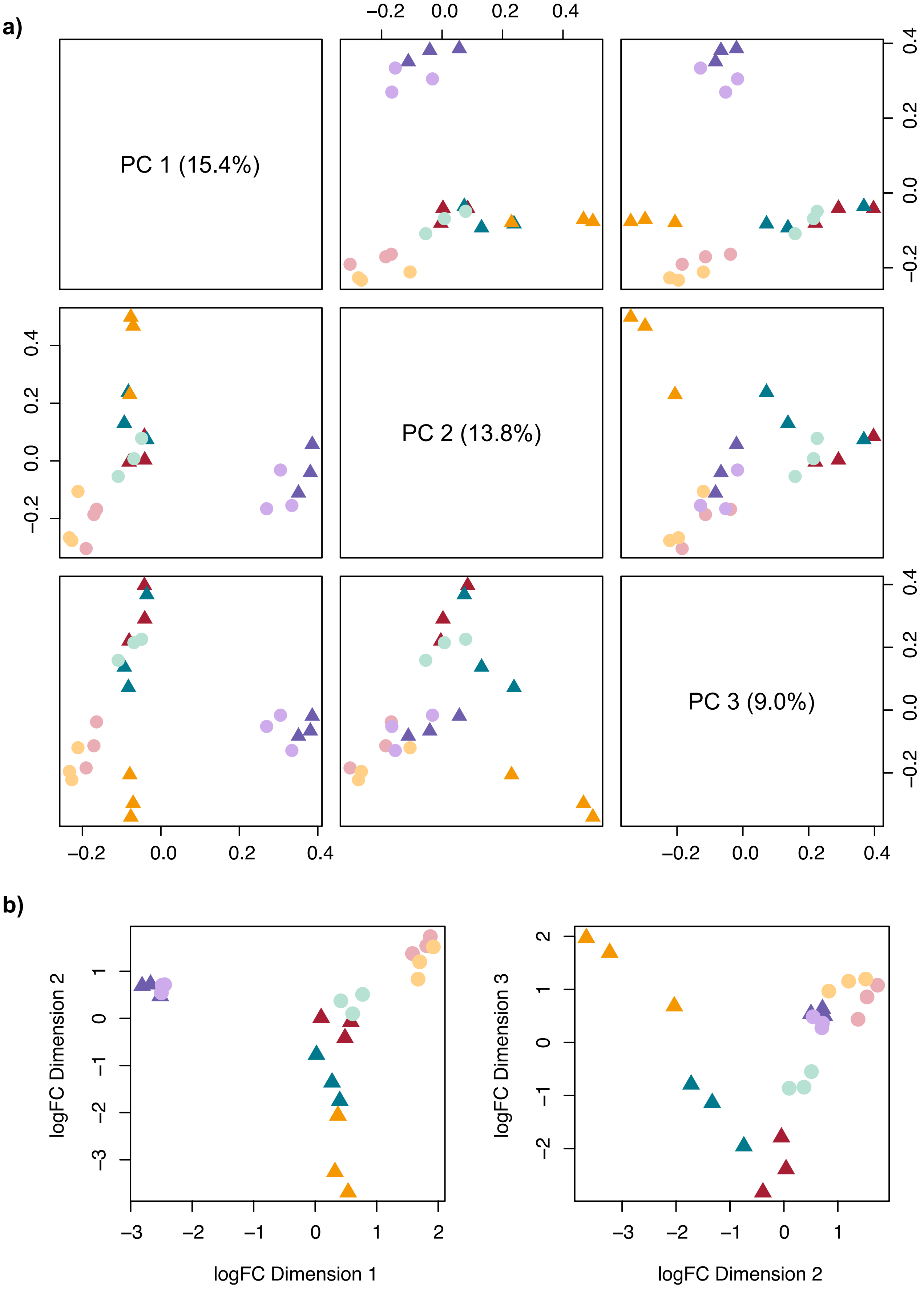

**Figure S6.** Boxplots showing the degree of gene expression divergence between populations of the alpine and montane ecotype, respectively. The difference between the means was statistically tested using Wilcoxon signed rank test and found to be not significant (p = 0.56).

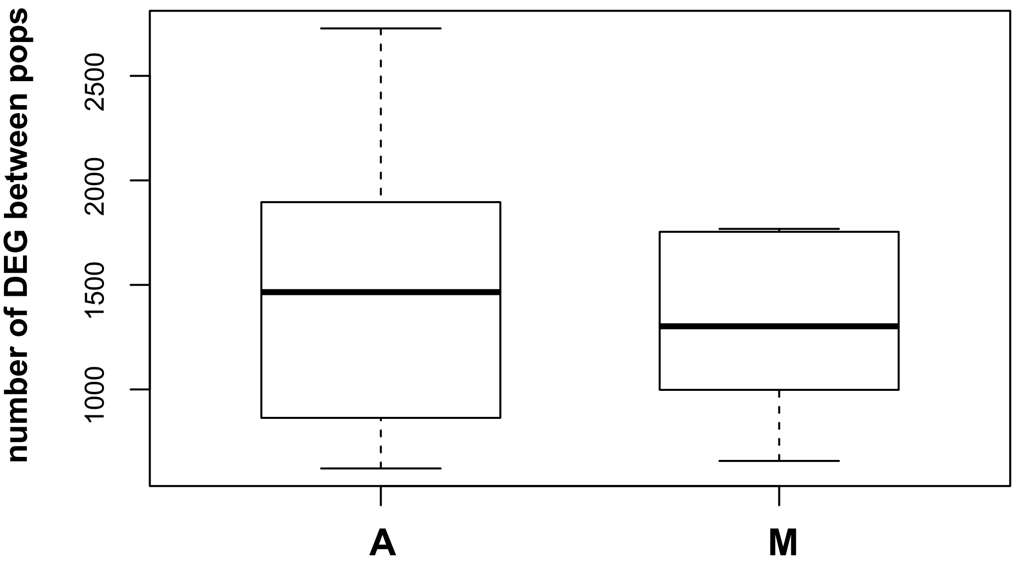


**Figure S7.** Z-score distribution obtained from the cRDA transcripts scores. Dashed and full red lines correspond to significance thresholds of p-value of 0.05 and 0.01, respectively.

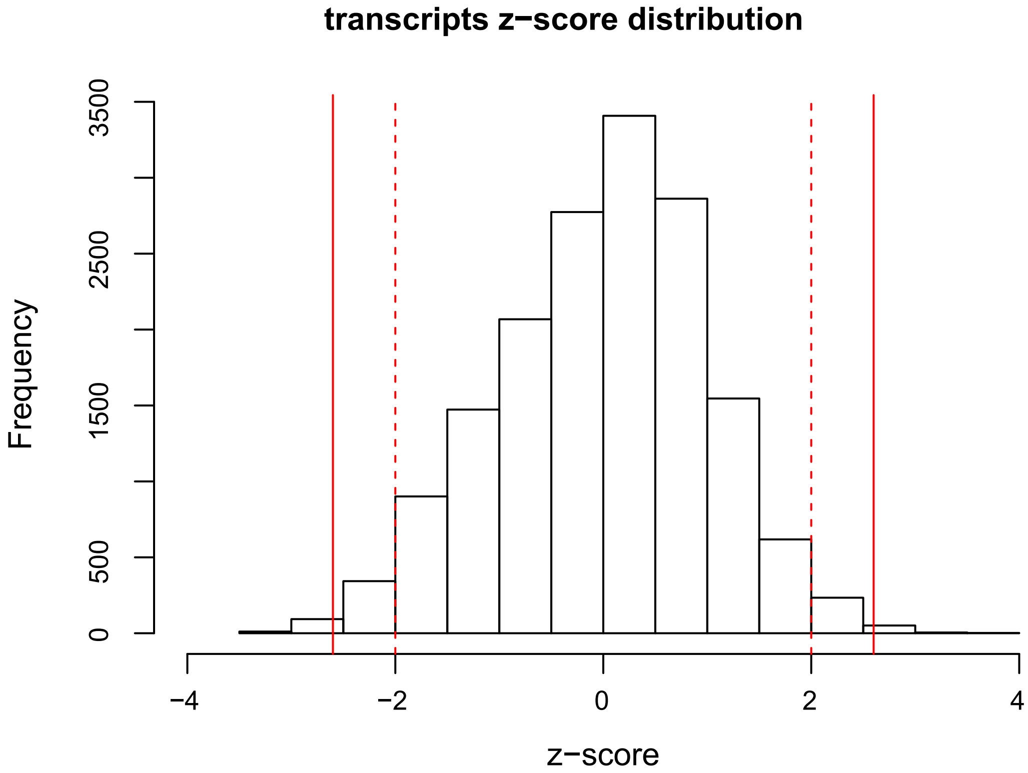

**Figure S8.** Venn diagrams showing overlaps between lists of DEGs and 739 cRDA outliers. Genes underexpressed (a) and overexpressed (b) in the montane ecotype compared to the alpine at each pair and in the cRDA analysis.

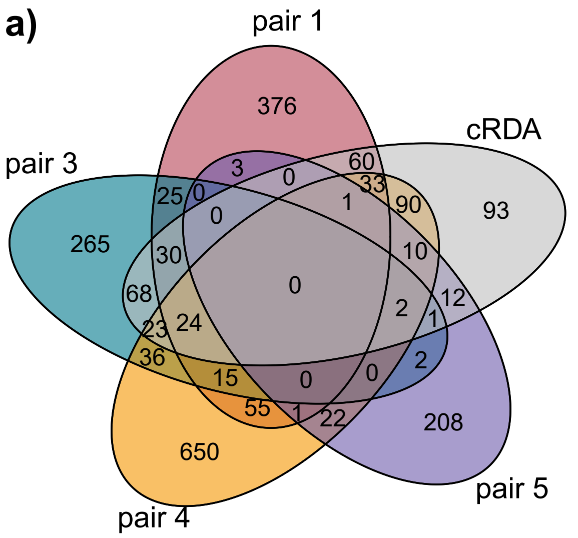


**Figure S9.** Distribution of $F_{ST}$ in 1,000 randomly selected genes (white bars) compared to DEGs (black bars) in ecotype pair 1 (**a**) and 3 (**b**). Dotted lines show the mean $F_{ST}$ values.

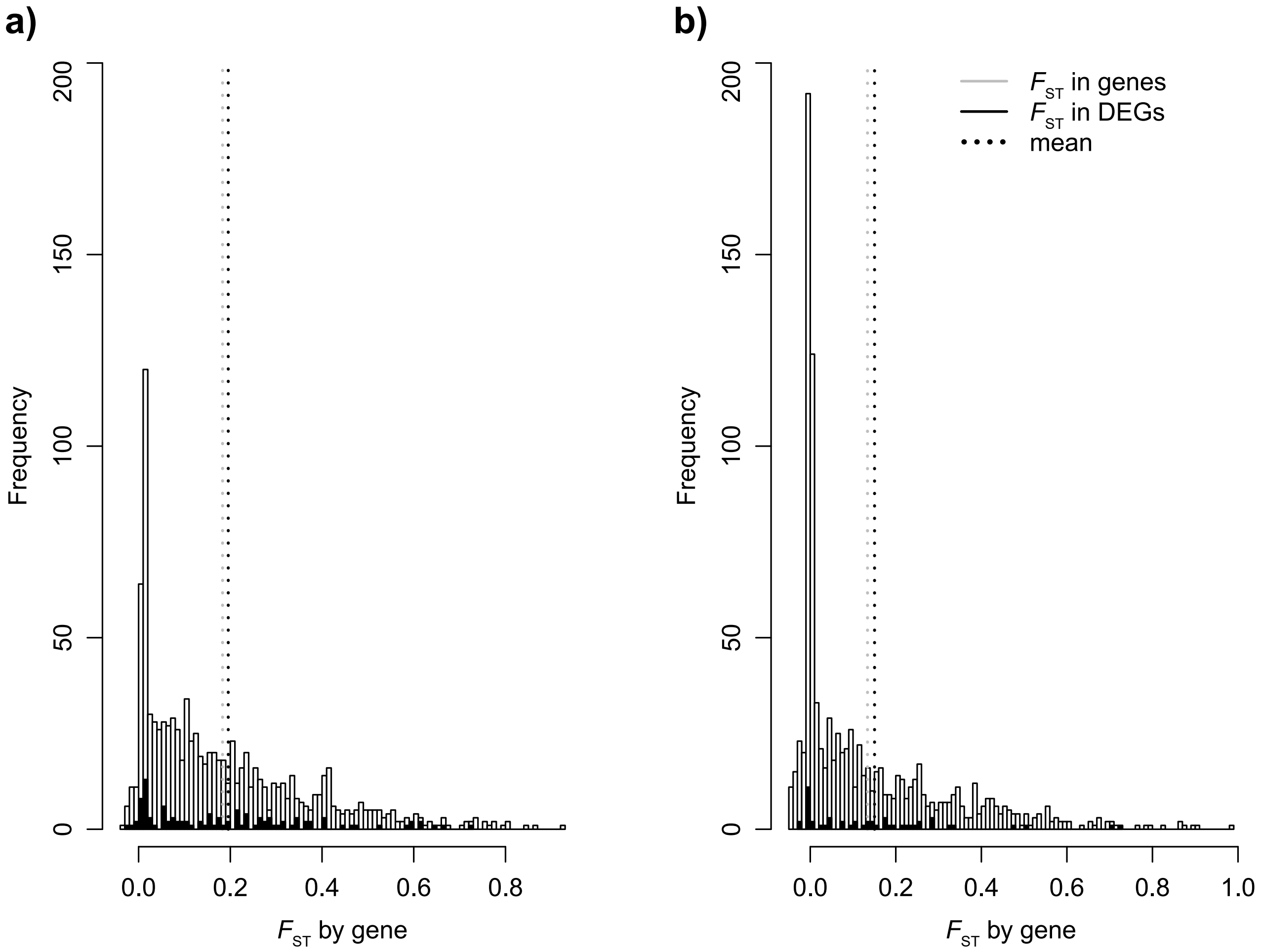
